# Integrated spatiotemporal transcriptomic resolution of embryonic palate osteogenesis

**DOI:** 10.1101/2023.03.30.534875

**Authors:** Jeremie Oliver Piña, Resmi Raju, Daniela M. Roth, Parna Chattaraj, Fahad Kidwai, Fabio R. Faucz, James Iben, Apratim Mitra, Kiersten Campbell, Gus Fridell, Caroline Esnault, Ryan K. Dale, Rena N. D’Souza

**Author notes:** Corresponding author., Chief, Section on Molecules & Therapies for Craniofacial & Dental Disorders, Eunice Kennedy Shriver National Institute of Child Health and Human Development National Institutes of Health 9000 Rockville Pike, Building 50, 4120 Bethesda, MD, USA 20892.

## Abstract

The differentiation of osteoblasts and the subsequent formation of bone marks an important terminal phase in palate formation that leads to the separation of the oral and nasal cavities. While the developmental events that precede palatal osteogenesis are well explored, major gaps remain in our understanding of the molecular mechanisms that lead to the bony union of fusing palatal shelves. Herein, the timeline of osteogenic transcriptional programming is unveiled in the embryonic palate by way of integrated bulk, single-cell, and spatially resolved RNA-seq analyses. We define spatially restricted expression patterns of key marker genes, both regulatory and structural, that are differentially expressed during palatal fusion, including the identification of several novel genes (*Deup1, Dynlrb2, Lrrc23*) spatially restricted in expression to the palate, providing a relevant framework for future studies that identify new candidate genes for cleft palate anomalies in humans as well as the timing of mammalian embryonic palatal osteogenesis.

## INTRODUCTION

Palatogenesis involves a complex interplay between cell adhesion molecules, epigenetic regulators and transcription factors that function within signaling pathways to coordinate the growth, migration, elevation, and fusion of cranial neural crest (CNC) cell derivatives^1, 2^. How these heterogeneous signaling networks define the morphogenetic gradients that govern patterning morphogenesis of anatomically distinct regions of the embryonic palate is not well understood. The development of the primary palate from the medial nasal processes occurs anterior to the incisive foramen and involves a set of genes that are distinct from those involved in the formation of the secondary palate. The latter is derived from the bilateral maxillary processes, that grow into the palatal shelves, posterior to the incisive foramen^2^. The secondary palate is further sub-divided into the anterior (hard, osseus) and posterior (soft, muscular) secondary palate. These anatomical regions that utilize different morphogenetic codes parallel the phenotypic heterogeneity of orofacial cleft conditions observed in humans^3^.

The dynamic morphogenetic processes required for complete development of the palate are often disrupted by genetic and/or environmental insults^3–6^. Palatal clefts (a sub-set of orofacial cleft anomalies) together with cleft lip are among the most common birth defects in humans, occurring in approximately 1 in 700 live births^7, 8^. Such birth defects pose significant physical, mental, psychosocial, and financial burden on patients and their caregivers throughout life, often requiring multiple stages of complex surgeries with varying rates of success^9, 10^. The pathophysiology of palatal clefts is complex and is proposed to involve disturbances in cell proliferation^11^, migration^12^, epithelial-to-mesenchymal transformation (EMT)^13^, cell-cell adhesion^14^, or terminal osteogenic differentiation^15^, leading to the failure of palatal shelf fusion^8^. It is known that osteogenesis is a key stage of palatal shelf in-growth and secondary palate fusion, the failure of which can result in a sub-mucous cleft palate, wherein a viable bony bridge separating oral and nasal cavities fails to form^16^. While the stages of palatogenesis have been studied and characterized morphologically, the basic molecular mechanisms driving embryonic palatal fusion and osteogenesis remain elusive^1, 17^. If properly explored, such information can be applied to the development of pre-clinical models to trial novel therapeutics that may benefit individuals with isolated or syndromic cleft palate defects who currently face complex surgeries and the arduous burden of life-long care without innovative solutions^9^.

To meet this critical translational need, we identified the spatiotemporal differential gene expression signatures of cell populations driving secondary palate fusion. Specifically, we observed the morphogenetic gradients of expression of key osteogenic cues driving the patterning of the secondary palate through time and space. Our studies sought to gain a global, whole-transcriptome view of the secondary palate during development by employing bulk-RNA-sequencing (RNA-seq) of total RNA from microdissected murine embryonic palatal shelves (E13.5-P0) (Fig. 1a). We then resolved specific cell populations at single-cell resolution by way of single-cell RNA-sequencing (scRNA-seq) of early embryonic stages leading to palatal fusion (E13.5-E15.5), which we define as the onset of the dissolution of the midline epithelial seam (Fig. 3a). To provide *in vivo* validation of select differential marker genes identified, multiplexed *in situ* hybridization (RNAScope) was then performed on the same stages of development. Finally, we contextualized whole-transcriptome gene expression patterns *in situ* at the stage of active osteogenic differentiation (as identified from the bulk and scRNA-seq datasets, E14.5-E15.5) by way of spatial RNA-seq (spRNA-seq, Visium, 10x Genomics), offering a first-of-its-kind spatiotemporal resolution of transcriptomic profiles of the developing secondary palate (Fig. 6a). Importantly, by providing this multimodal transcriptome-wide dataset to the craniofacial biology community through open-access sharing of all data on FaceBase (the primary shared data source for craniofacial researchers worldwide), we aim to support and facilitate translational studies on normal and abnormal palate formation.

**Fig. 1.**
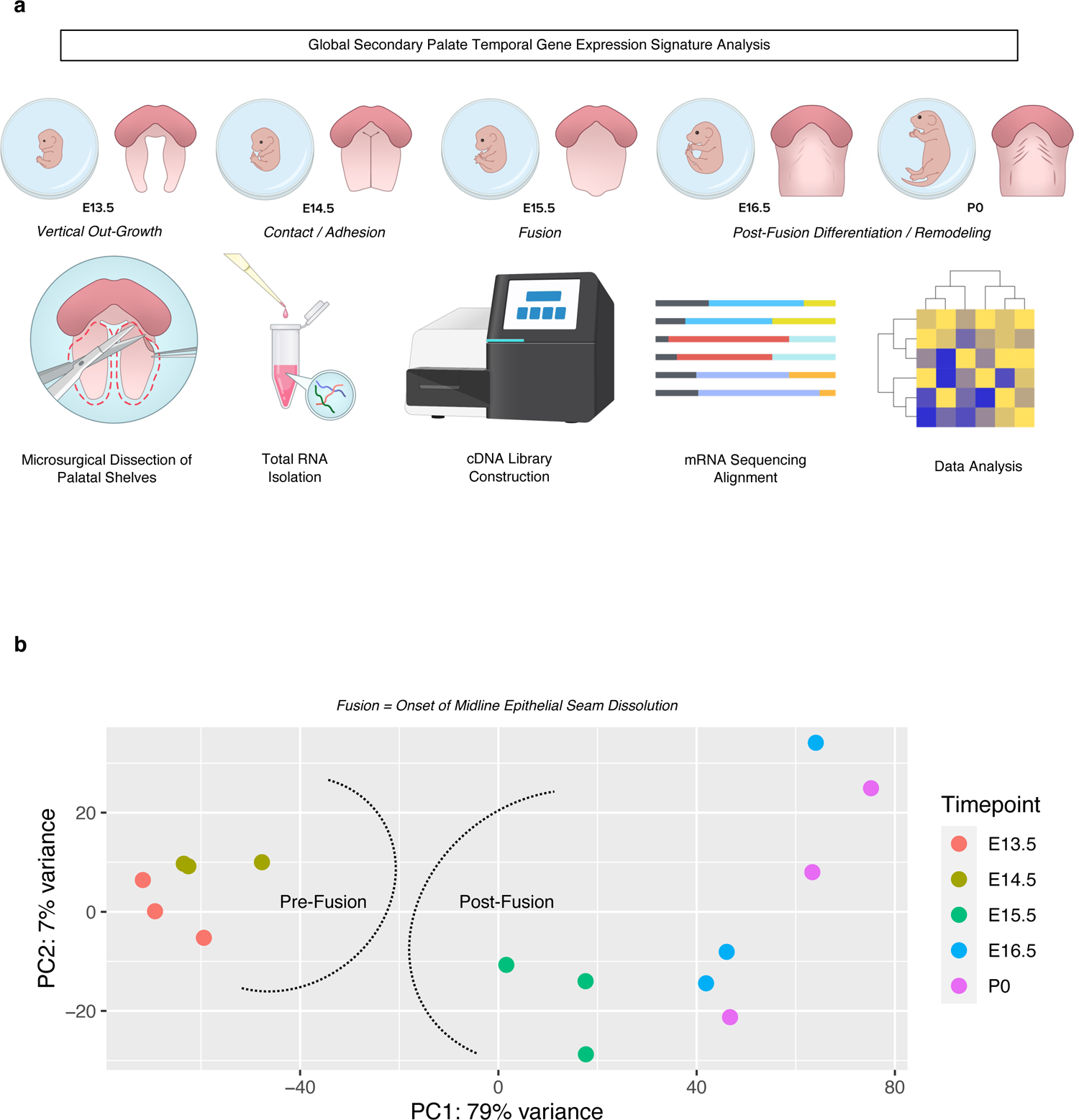
Global transcriptomic profiling of the embryonic secondary palate reveals unique signatures pre- and post-fusion. **a** Total RNA was isolated from microdissected palatal shelves from embryonic stages E13.5, E14.5, E15.5, E16.5, and P0 (n=3 pooled biological replicates per stage), then run through bulk RNA-sequencing. **b** Principal Component Analysis (PCA) of all replicates demonstrating global associations in earlier development (pre-fusion) vs later development (post-fusion) of secondary palate.

## RESULTS

### Bulk RNA-sequencing captures stage-specific global gene expression signatures during palate development

Transcriptomic signatures were broadly grouped into pre-fusion (E13.5-E14.5) and post-fusion (E15.5-P0) clusters, as we defined the fusion event as the onset of midline epithelial seam dissolution which is known to occur between E14.5-E15.5^1^ (Fig. 1b). Our data suggest that a distinct transcriptomic shift may exist bordering the developmental milestone of palatal fusion. Furthermore, through differential comparisons of adjacent stages of development, we identified the stage of palatal fusion to demonstrate the greatest overall shift in gene expression based on Z-score (gene abundance) profiling and differential expression (Supp. Fig. 1). Thus, we hypothesized that this transitory stage of development, based on its genetic expression profile, likely comprises molecular genetic events of morphologic importance during palate development.

### Palatal osteogenesis begins with fusion

Through differential expression analysis of adjacent stages, we noted the onset of osteogenic programming by E15.5, supported by numerous marker genes that either drive or mark osteoblast commitment and differentiation (*Bglap/2/3, Mmp13, Dmp1, Spp1, Ibsp, Bmp8a, and Alpl*) as well as those that mark the formation of critical structural components of bone (*Col1a1, Col1a2, Sparc*) differentially up-regulated compared to E14.5, as well as negative regulators of Msx1 activity (*Msx3*), BMP signaling (*Skor1*), neural crest migration (*Gbx2*), and skeletal matrix differentiation (*Hoxc5*) down-regulated at the same stage (Fig. 2a, Supp. Fig. 1). This onset of osteogenic programming was corroborated by functional enrichment analysis on differentially expressed genes at the stage of palatal shelf fusion (E14.5-E15.5), highlighting genes and pathways associated with osteoblast differentiation and function (Supp. Fig. 2). Notably, gene markers indicative of osteoblast commitment and differentiation universally increased in expression *only* from E14.5-E15.5. Expression patterns of key up-stream regulatory transcription factors known to orchestrate palatogenesis, including *Twist1*, *Twist2*, *Barx1*, *Msx1*, *Tbx22*, *Meox2*, and *Pax9* (Fig. 2b) support the osteogenic switch in transcriptomic signature occurring at this stage of development. Global transcriptomic signatures in pre-fusion versus post-fusion embryonic palate tissue led us to focus our analysis on stages that precede osteogenesis and those that drive bony fusion at higher resolution by way of scRNA-seq and spRNA-seq modalities.

**Fig. 2.**
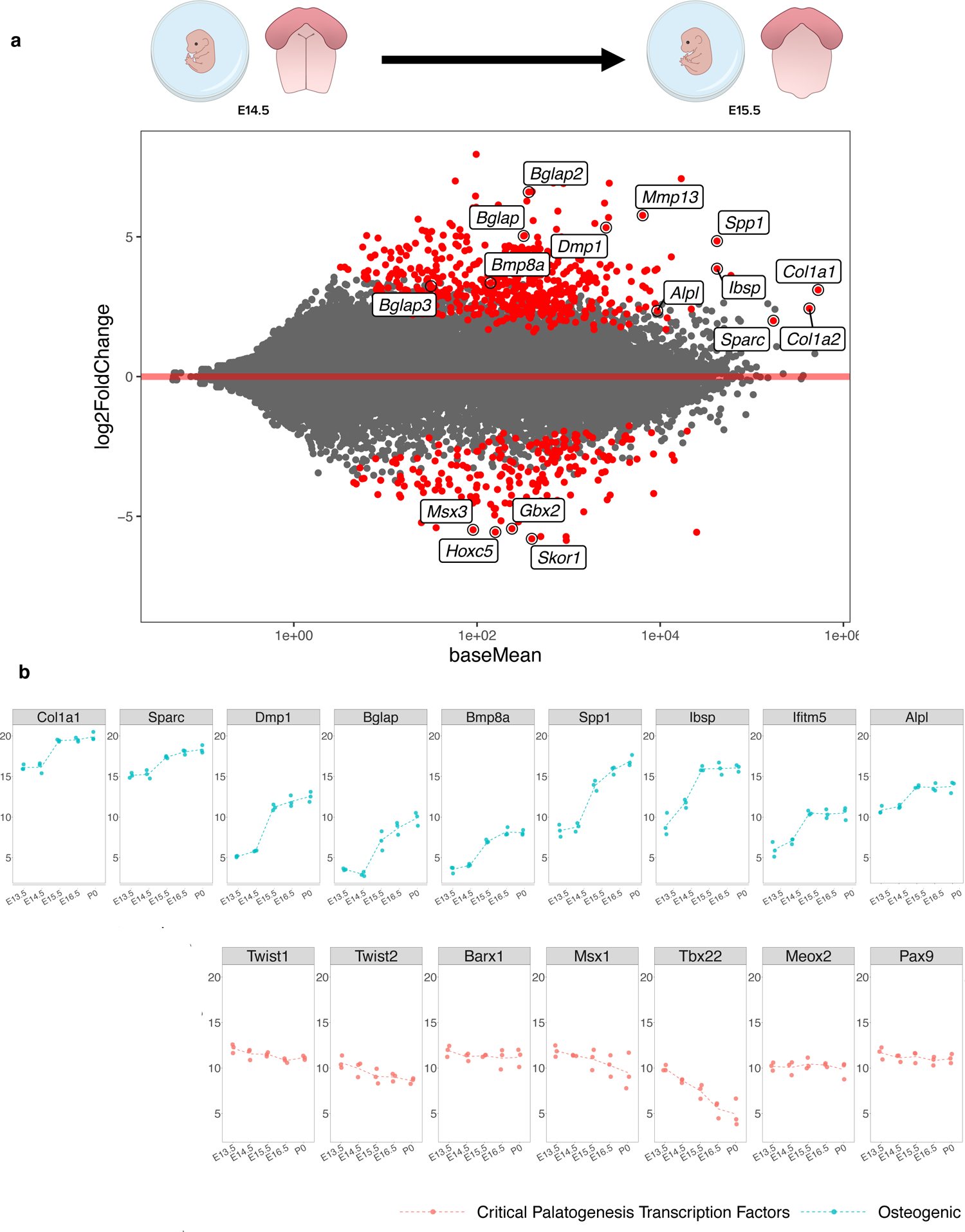
Differential shift toward osteoblast commitment defines the transition from E14.5 – E15.5 in palate development. **a** M-A plot highlights statistically significant (red) differentially expressed genes identified in transcriptomic comparison of E14.5 to E15.5 in development. Genes related to osteoblast lineage commitment and differentiation (up-regulated: *Bglap/2/3, Col1a1, Col1a2, Dmp1, Spp1, Alpl, Sparc, Mmp13, Bmp8a*; down-regulated: *Msx3, Gbx2, Hoxc5, Skor1*) were only shown to be differentially expressed at this transitory stage of palatal fusion. b Plotting individual genes across all time points studied demonstrated the noted shift in relative osteogenic gene abundance from E14.5 – E15.5, while the same shift was not seen across known critical transcription factors involved in palatogenesis, such as Twist1/2, Barx1, Msx1, Tbx22, Meox1, and Pax9.

### Single-cell RNA-seq reveals heterogeneity of palatal shelf cell populations and corroborates osteogenic marker abundance in E14.5-E15.5 transition

To further dissect the individual cell populations driving the morphogenetic changes contributing to palatal shelf fusion, droplet-based scRNA-seq was performed on pools of age-matched littermates’ microdissected palatal shelves at E13.5, E14.5, and E15.5 (Fig. 3a). Integration and clustering using supervised parameter selection revealed 11 unique cell populations which were shown to be heterogeneous across development stages (Fig. 3b). Osteogenic markers were identified to be associated primarily with two clusters, Clusters 1 and 3. Markers of Cluster 1 included primarily osteoblast markers, including *Col1a1*, *Col3a1*, and *Postn*, while Cluster 3 markers included genes of pre-osteoblast affinity (*Runx2*, *Sox11*, as well as *Col1a1*) (Fig. 3c-d). Markedly, the regulatory transcription factor orchestrator of osteogenic commitment, *Runx2*, demonstrated earlier stage and greater enrichment of expression compared to the structural component marker of bone, *Col1a1* (Fig. 3d). These clusters show differential increase through development, with other markers of osteoblast commitment appearing in later stages, including *Sparc*, *Alpl*, and *Bmp6*. Notably, the relative *in vivo* cell type abundances might not be reflective of cluster proportions due to cell isolation biases and strategies.

**Fig. 3.**
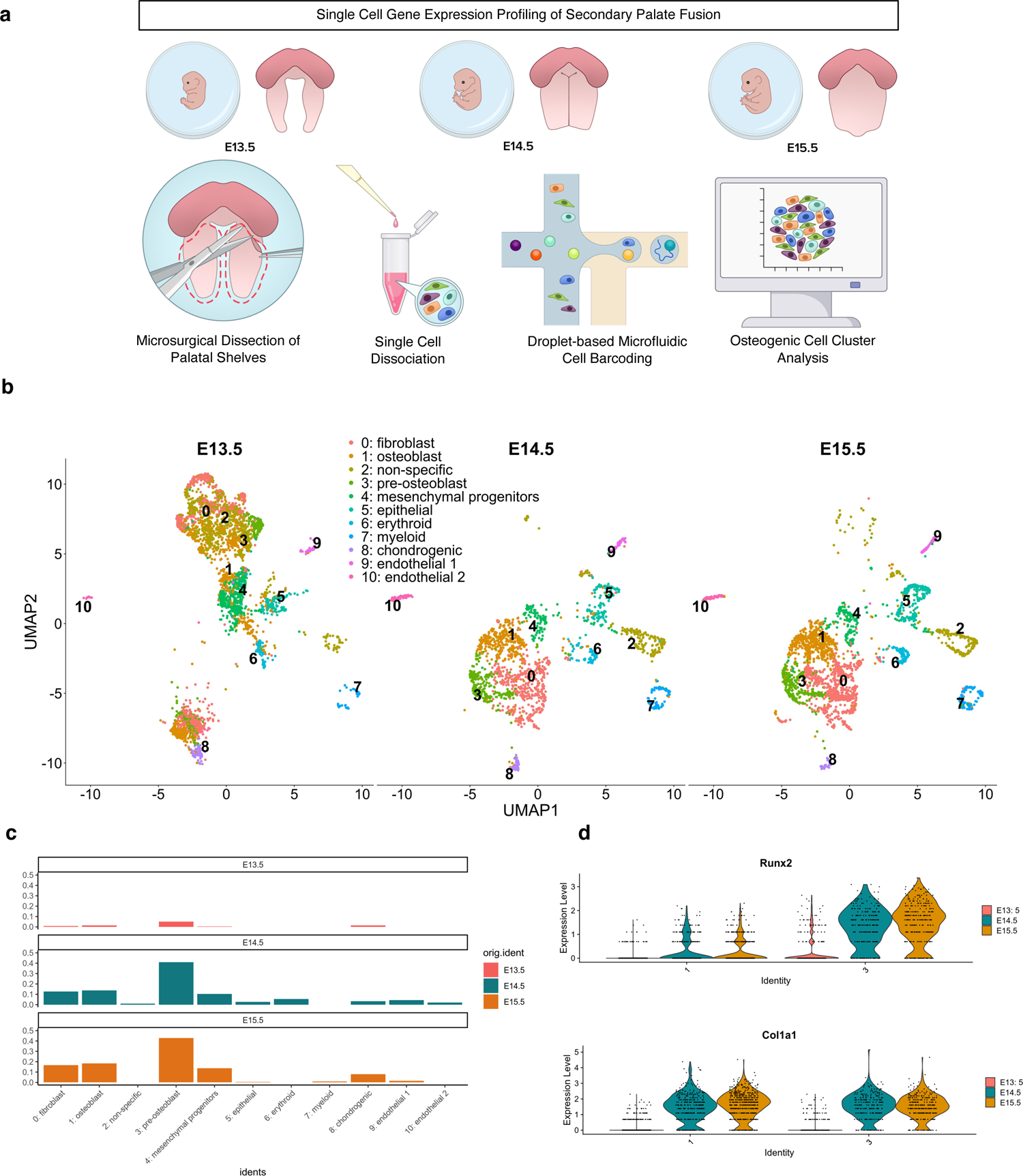
Single-cell-level transcriptomic profiling of the embryonic secondary palate. **a** Microsurgical dissection of secondary palatal shelves in E13.5, E14.5, and E15.5 mouse embryos was performed to isolate single cell suspensions for droplet-based single-cell RNA-sequencing (10x Genomics). **b** Staged clusters were integrated with Seurat, with noted cell type and temporal heterogeneity; Known markers of specific cell populations were associated to defined clusters from integrated dataset. Each cluster was labeled distinct cell types based on known functional gene markers, as follows: 0: Fbn2, Col23a1, Mmp16; 1: Col1a1, Col3a1, Postn; 2: Sdk1, Atp8b4; 3: Runx2, Col1a1, Sox11; 4: Kif11, Gpc3, Prr16; 5: Cldn7, Epcam, Krt18; 6: Alas2, Gypa, Hbb-bt; 7: Laptm5, Tyrobp, Dock2; 8: Col2a1, Col9a1, Matn1; 9: Cdh19, Ednrb, Egfl8; 10: Emcn, Kdr, Cdh5. **c** Histogram split by timepoint depicting relative proportions of cells from all 11 clusters. **d** Violin plots of individual genes denoting increased expression of pre-osteoblast and osteoblast commitment markers by E14.5-E15.5.

We observed the most qualitative differences in the scRNA-seq UMAP plots between E13.5 and E14.5, while the bulk RNA-seq differential expression analyses highlighted a greater difference between E14.5 and E15.5. We reasoned that this discrepancy might be due to cell type-specific changes picked up in scRNA-seq that were masking each other in bulk RNA-seq. However, pseudo-bulk analysis of the scRNA-seq data was still consistent with an observed transition between E13.5 and E14.5. We therefore hypothesize a sharp, transient transition in expression profile that occurs around E14.5 that our one-day sampling resolution is currently unable to resolve. While the timepoints may not be perfectly aligned across the assays due to differing pregnancy starting time points in hours, we nevertheless observe upregulation of genes at later timepoints compared to earlier timepoints that are consistent between assays.

### *In vivo* validation and spatiotemporal mapping of select osteoblast markers in palatal shelves during the transitory fusion stages

To interrogate the *in situ* transcriptomic profile of the verified osteoblast markers in the secondary palate during fusion, we employed multiplexed *in situ* mRNA hybridization (RNAScope) of select marker genes (*Runx2*, *Col1a1*, *Sparc*, *Sost*). Importantly, we sought to co-localize across space and time marker gene transcripts indicative of 1) regulatory activation of osteogenic differentiation (*Runx2*, *Sost*) and 2) structural components of newly forming bone tissue (*Col1a1*, *Sparc*) *in situ*, noting the spatiotemporal interplay between these stages of osteogenic programming. *Runx2* is known as a reliable marker for pre-osteoblasts^18^, *Sparc* for functional / mature osteoblasts (required for successful osteoblast mineralization of osteoid)^19, 20^, *Col1a1* for functional / active osteoblasts^21, 22^ (while not exclusive to osteoblasts, it is closely associated in its genetic interaction to *Runx2* and *Sparc* per String DB [Supp. Fig. 3] – also providing a useful co-expressing *in vivo* perspective in surrounding matrix tissues of the craniofacial complex), and *Sost* for Wnt-regulatory osteocyte specificity^23, 24^. This allowed for *in vivo* resolution of osteogenic cell relationships identified from the integrated bulk and scRNA-seq workflows as regulatory mechanisms of osteogenic induction in turn fuel the transcriptional programming for structural components of bone morphogenesis (Fig. 4a). Comparative mid-palatal formalin fixed paraffin embedded (FFPE) coronal sections from E13.5, E14.5, and E15.5 embryonic stages were selected based on craniofacial complex anatomical landmarks (bulbous nasal septum, tongue attachment visible). We observed spatiotemporal expression patterns corroborating the sequencing studies, identifying greater density of expression in palatal shelf ossification centers beginning at E14.5 and increasing notably at E15.5. The expression of the pre-osteoblast marker, *Runx2*, localized throughout the developing palatal mesenchyme, with noted diffuse signal in central ossification centers. In contrast, the highly specific osteocyte marker, *Sost*, localized with much lower signal density in early stages of development (E13.5) – in line with RNA-seq quantification – with expression signal appearing selectively in E14.5, followed by a notable increase by E15.5 following palatal fusion (Fig. 4b). These spatiotemporal expression patterns may indicate distinct functional roles for osteoblasts along their developmental trajectory during palatal mesenchyme proliferation, differentiation, and fusion.

**Fig. 4.**
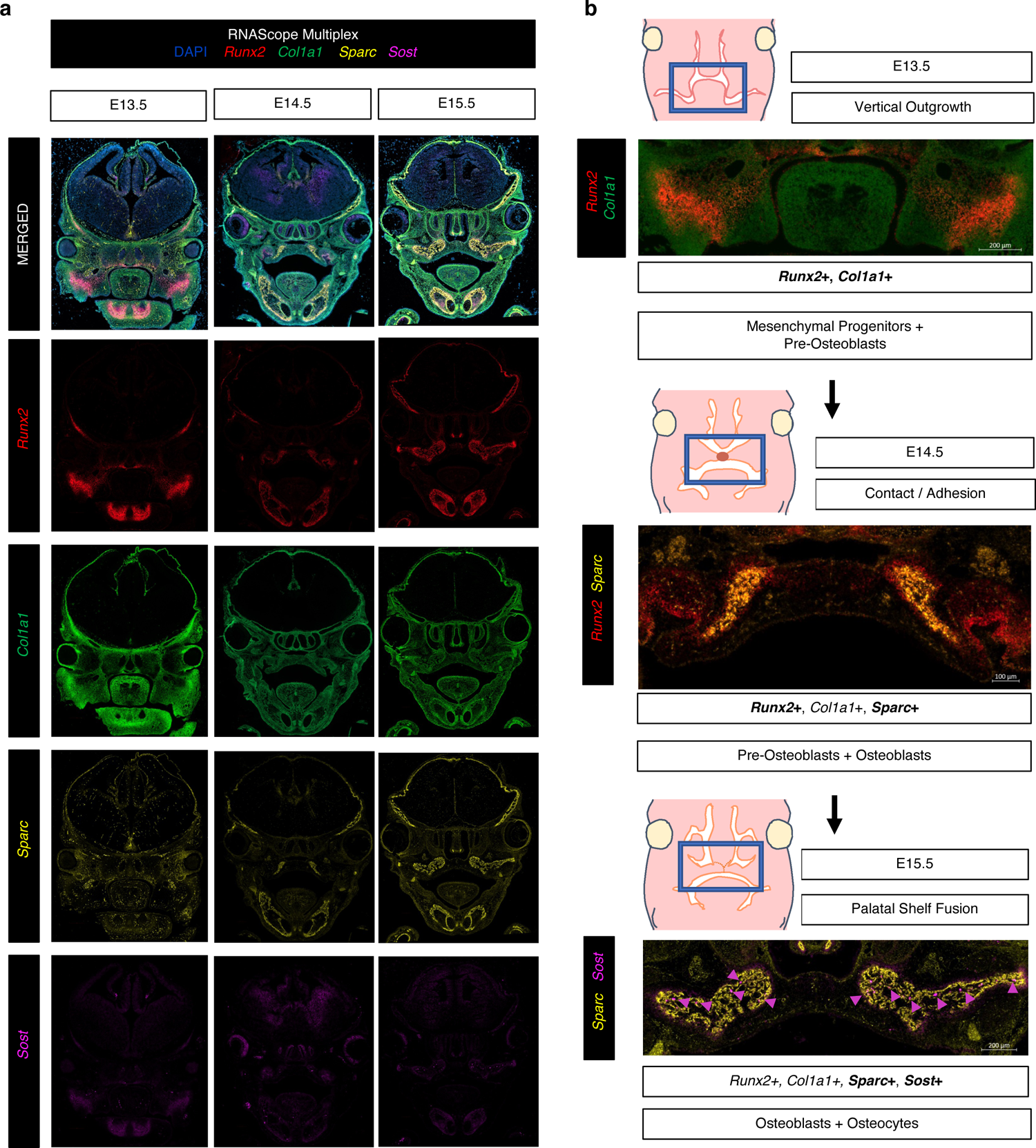
RNAScope Multiplex for in vivo validation of spatiotemporal osteoblast marker mRNA expression in secondary palate tissues. **a** Select osteogenic marker genes hybridized to track lineage-specific spatial patterns of differentiation: Runx2 (pre-osteoblast), Col1a1 (early functional osteoblast), Sparc (mature functional osteoblast), and Sost (osteocyte), in mid-palatal coronal sections via RNAScope Multiplex (*All in vivo validation experiments performed in biological and technical triplicate). **b** Proposed spatiotemporal osteogenic cell lineage activation sequence in developing secondary palate mesenchyme.

### *In situ* spatial RNA-seq of palatal fusion enables high-resolution *in vivo* context to palatal mesenchymal differentiation profiles

While bulk and scRNA-seq approaches to study the developing secondary palate provide meaningful global temporal transcriptomic data, these approaches lack tissue-specific spatial information critical to inferring genetic regulatory networks *in vivo*. Although some *in vivo* validation can occur through the lens of RNAScope, it is primarily a qualitative assay. Further, these *in vivo* marker validation assays are inherently limited due to multiplexed channel maximums within a narrow spectrum of visibility. This is of particular importance in determining the dynamic nature of the developing secondary palate, from proliferating, migrating CNC cell progenitors to functionally differentiated cell players. Moreover, it is known^25^ that the single-cell suspension, dissociation, and fixation processing steps involved in the 10x Genomics scRNA-seq workflow carry an inherent risk of altering cellular homeostasis. This could lead to physiologically unimportant differential expression observed in cell clusters. We therefore employed spRNA-seq (Visium, 10x Genomics) on FFPE coronal mid-palatal sections from E14.5 and E15.5 embryos to enable real-time assessment of *in situ* gene expression and generate a more complete picture of secondary palate fusion (Fig. 5a). While the per-cell resolution is lower than the scRNA-seq workflow, the spatial biological insight from sequencing relatively undisturbed tissue slices *in situ* brings higher confidence in observed spatiotemporal gene expression trends.

**Fig. 5.**
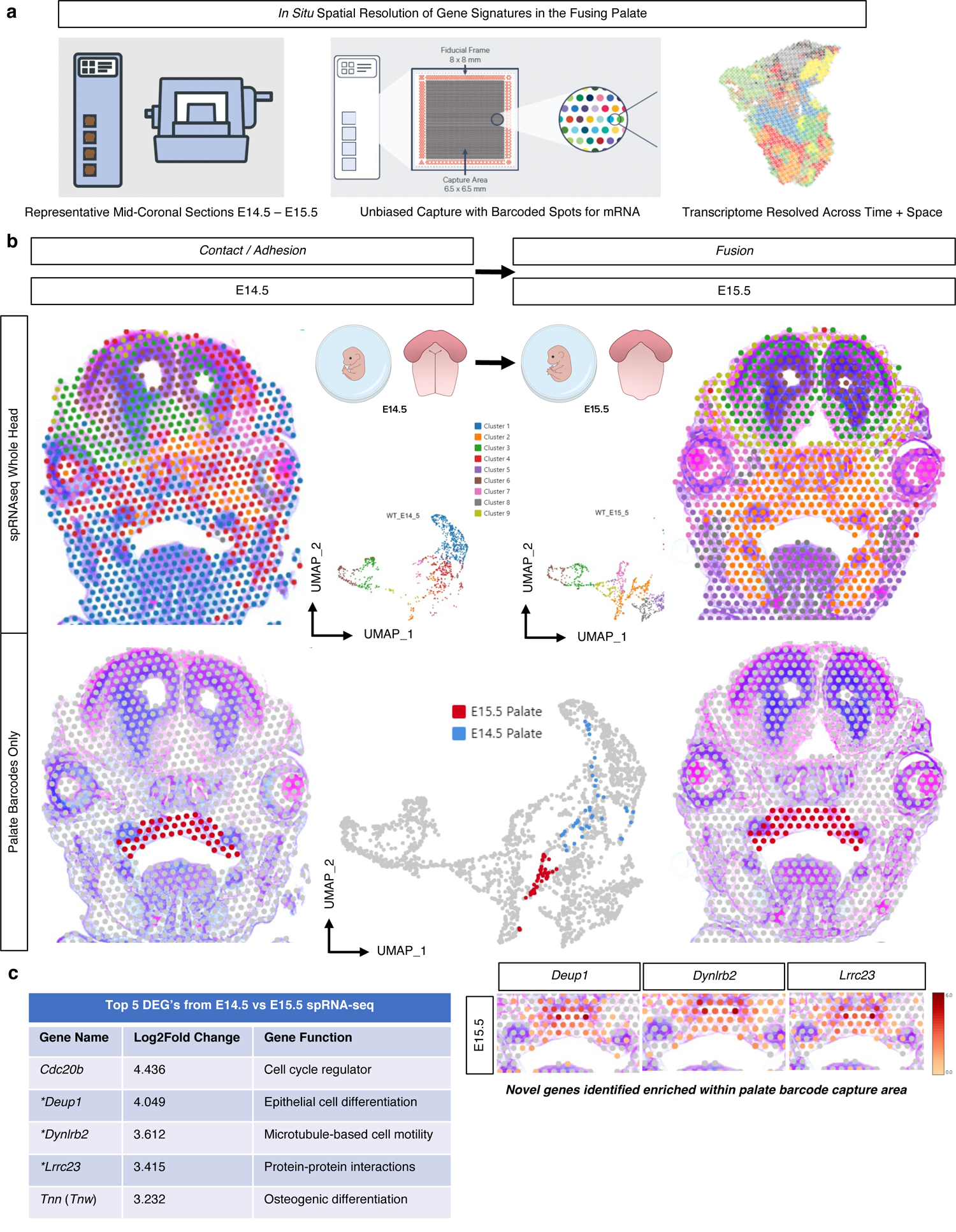
Spatial RNA sequencing (spRNA-seq) for the in situ resolution of the fusing palate’s transcriptome. **a** Mid-palatal coronal cross sections of whole embryo heads were placed on barcoded Visium slide. **b** *In vivo* clusters were defined from whole embryo head, demonstrating spatial relationships and morphogenetic diversity of expression, further filtered for only those clusters encoded on the barcodes placed within the palate tissue in each respective section to identify top differentially expressed genes (DEG’s) **c** from E14.5 vs E15.5 in the palate. *Denotes novel palate enriched genes identified in spRNA-seq.

Spatial cell cluster heterogeneity was observed globally between distinct morphological zones of the craniofacial complex in coronally-oriented mid-cranial sections from E14.5 and E15.5 embryonic stages (Fig. 5b). Barcodes representing the secondary palate *in situ* tissue arrangement were manually selected within the 10X Genomics Loupe Browser, allowing for a targeted integration of transcriptomic data specific to the palate in each section. Whole-transcriptome differential expression of selected palate tissue barcodes between E14.5 and E15.5 tissue sections highlighted marker genes of potential significance in the processes of secondary palate fusion, including genes related to cell cycle regulation, cilia functionality, cell motility, protein-protein interactions, as well as epithelial and osteogenic cell differentiation (Fig. 5c). To interrogate the spatiotemporal osteogenic cell lineage differentiation milestones in palatal mesenchymal cells at the time of fusion *in situ*, known markers of osteogenic progenitors (*Six2, Erg, Bmp7*), pre-osteoblasts (*Runx2*, *Sp7*, *Mmp13*), osteoblasts (*Ibsp*, *Ifitm5*, and *Spp1*), and osteocytes (*Sost*, *Dmp1*, *Phex*) were identified spatially at each stage (Supp. Fig. 4a-d). Markers from early osteogenic cell types (osteoprogenitors, pre-osteoblasts, and osteoblasts) demonstrated some expression at E14.5; however, all osteocyte conserved markers only appeared in substantial expression by E15.5 – remaining spatially restricted to the mineralized bone (lateral), with some condensed expression noted within the midline site of fusion (mesial), perhaps suggesting multiple spatially distinct regions of osteogenic differentiation within the fusing palate mesenchyme.

### Spatial characterization of fusing palatal shelves *in situ* unveils novel marker genes

With an eye toward discovery, we turned our attention to the most highly differentially expressed genes identified from the spRNA-seq analysis to identify potential novel markers of palatal fusion across space and time (Fig. 5c). By selecting only for barcodes placed within the palate tissue catchment area on the Visium slide, this enabled us to directly compare whole-transcriptome spatiotemporal palate tissue-level gene expression changes, as the barcoded slide sequenced all mRNA transcripts bound to the oligonucleotide probes from the reference mouse transcriptome. The top five most highly differentially expressed genes (DEGs) from E14.5 – E15.5 were each up-regulated significantly, suggesting a potential functional role in propagating the necessary signaling cascades to promote palate cell differentiation and interaction at fusion. These top five DEG markers identified through spRNA-seq of palatal fusion were then compared to our previously generated bulk RNA-seq dataset to corroborate temporal evolution of expression levels through all embryonic time points studied (Supp. Fig. 5b), which affirmed the differential up-regulation in expression specifically from E14.5 – E15.5. This suggests that these gene markers – some (*Tnn*^26–30^, *Cdc20b*^31^) well-established in the prior literature to play a pivotal role in palate development, and others (*Deup1*, *Dynlrb2*, *Lrrc23*) not previously cited – may in fact be notable targets for further investigation.

### Multimodal whole transcriptome integration

To integrate these multiple whole-transcriptome assays, we took a cross-section through the bulk, scRNA-seq, and spRNA-seq datasets to look closely at specific osteogenic cell type markers, *Runx2* (pre-osteoblast)^32^, *Alpl* (osteoblast)^33^, and *Phex* (osteocyte)^34^. Osteoblast lineage differentiation significantly increases within the secondary palate mesenchyme from E14.5-E15.5 (Fig. 6a-c). This suggests that by E15.5, intramembranous ossification of the secondary palate mesenchymal primordia is actively commencing, but most of these mesenchymal osteogenic lineage-committed cells have not yet reached terminal differentiation (osteocyte). Osteocytes likely become more functionally active in the palate mesenchyme during later stages of development following mineralization and maturation of bone matrix tissues.^35, 36^ This staged activation of specific osteogenic regulators may be of functional developmental importance to the palatal shelves, as overlapping signaling networks are tightly coordinating proliferation, migration (mesial in-growth originating from lateral palatal ossification zones toward the mid-palatal suture mesenchyme), and terminal differentiation of pluripotent primordial palate mesenchymal cells (both lateral and mesial).

**Fig. 6.**
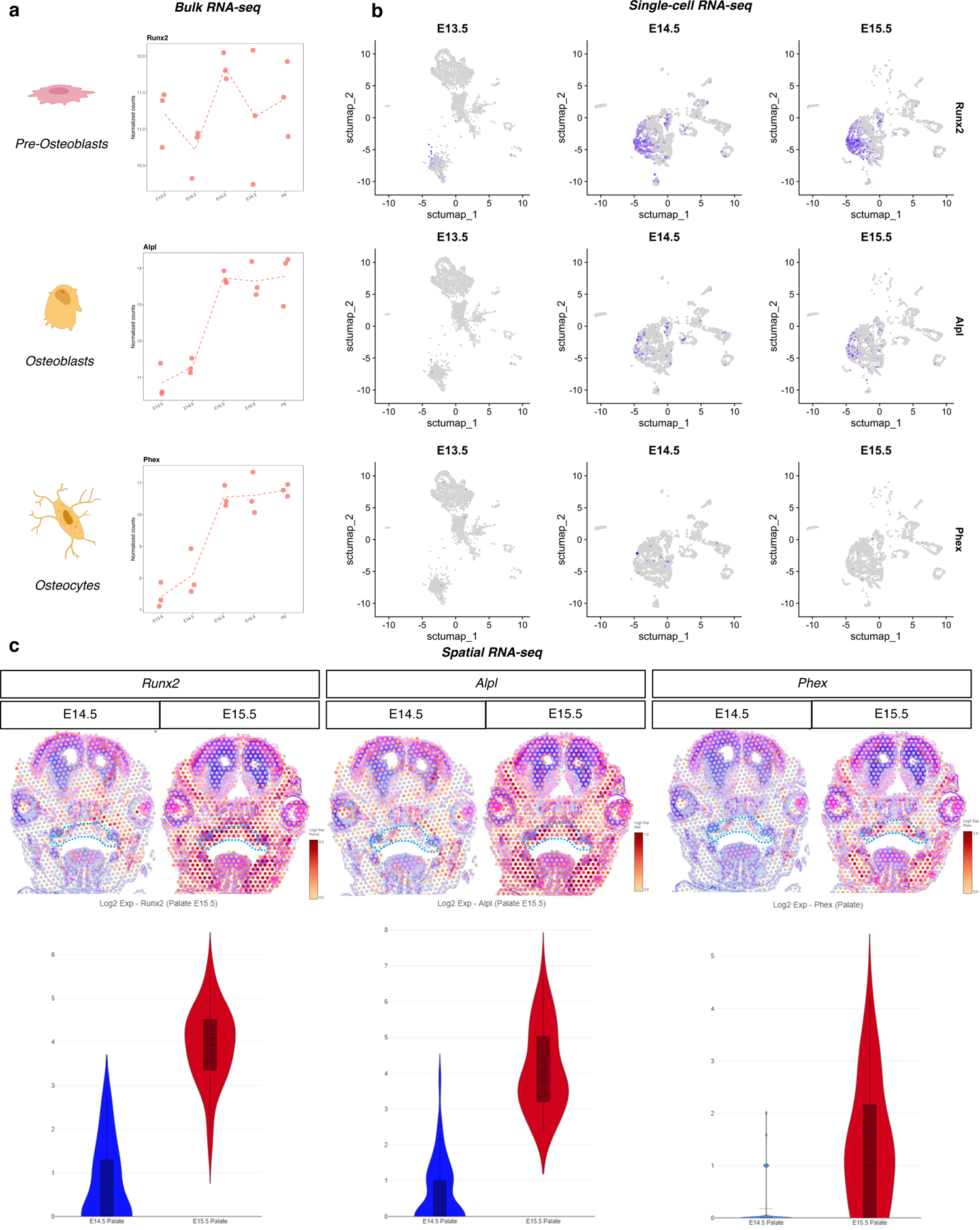
Integration of bulk, single-cell, and spatial transcriptomics. Bulk (a), single-cell (b), and spatial (c) resolution of select osteogenic differentiation markers delineates timeline of palatal osteogenesis and further characterizes the transcriptomic shift through contact, adhesion, and fusion of palatal shelves. *Blue dotted selection represents region of palate barcode selection (as shown in Fig. 5b) for spatial differential gene expression analysis.

**Fig. 7.**
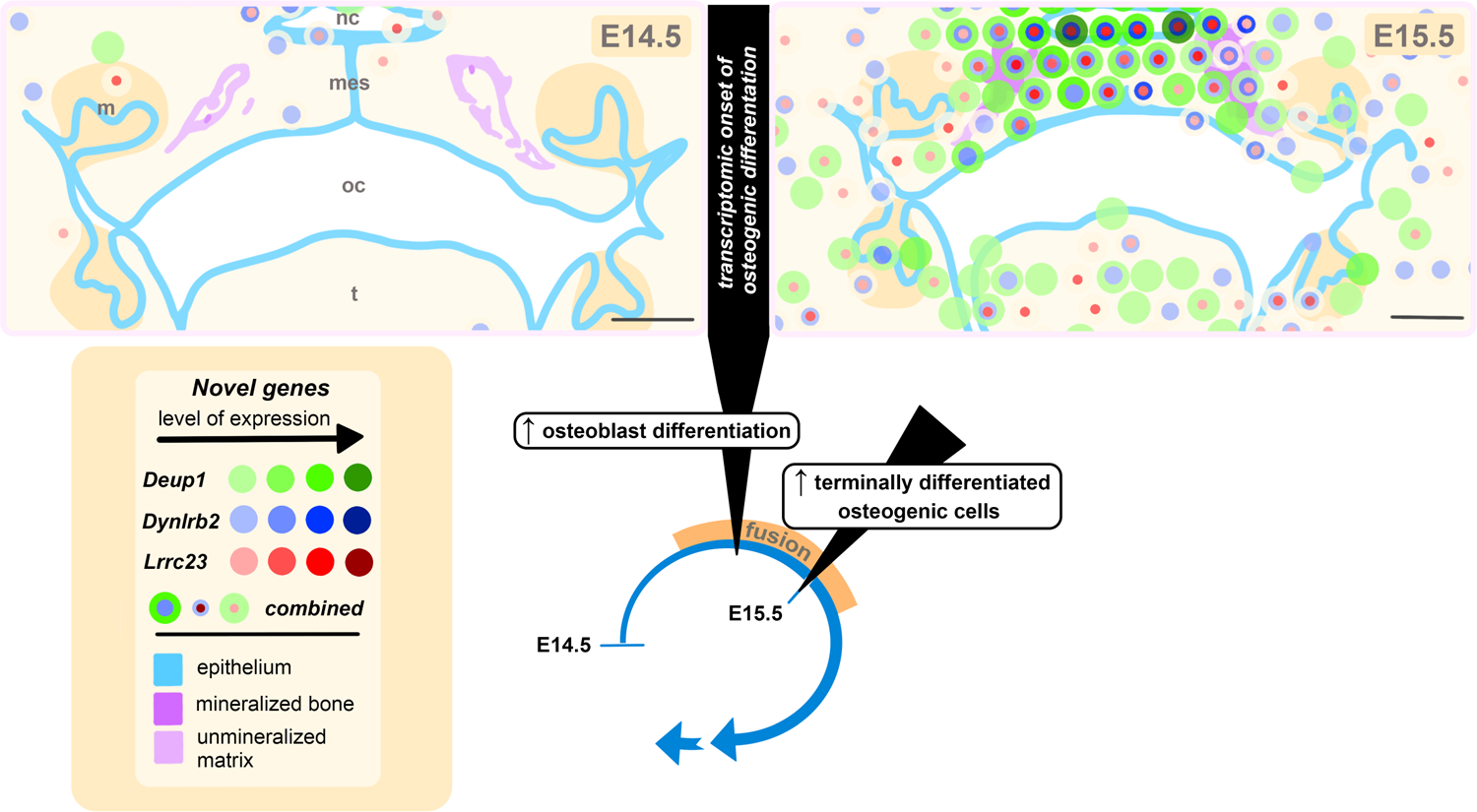
Spatiotemporal illustration of novel genes identified during palate fusion at the transcriptional onset of osteogenic differentiation. Graphical representation of murine secondary palate anatomy based on histological staining and multi-modal transcriptomic analysis identifying the onset of osteogenesis. This illustration highlights the epithelium, mineralized bone, and presumptive unmineralized matrix based on visual histological assessment. Dissolution of the midline epithelial seam (*mes*) marks the onset of palatal shelf fusion. During this period, there is a transcriptomic enrichment of markers of osteoblast differentiation, and an increase in terminally differentiated osteogenic cells at embryonic day 15.5 (E15.5). Colored circles correspond to 55 µm-diameter Visium transcriptomic resolution, demonstrating the expression patterns of three novel genes previously not described in the context of palate fusion. Increased expression levels, delineated using the 10X Genomics Loupe Browser, are represented here with darker shades of green (*Deup1*), blue (*Dynlrb2*), or red (*Lrrc23*). The combined localization of these novel genes is indicated by overlapping concentric colored circles, the diameter of which does not correspond to degree of expression. *nc*: nasal cavity, *oc*: oral cavity, *t*: tongue, *mes*: midline epithelial seam; scale bar: 200µm.

## DISCUSSION

This spatiotemporal analysis of secondary palate development provides a transcriptome-wide, staged developmental template of morphogenetic gradients driving the fusion event. The timing of morphogenetic gradients controlling palatogenesis revealed here by whole-transcriptome profiling and targeted imaging methods provides a roadmap for the identification of essential effector molecules in more targeted studies. Such knowledge will enhance our understanding of the basic mechanisms involved in the proper fusion of the palate, particularly the process of palatal osteogenesis, and can further be translated to human cleft palate conditions. Furthermore, key downstream effectors – now mapped through normal palatogenesis – can be examined further in murine disease models of cleft palate.

In our analyses, a transitory phase was identified between E14.5 (defined previously as the stage of contact/adhesion of palatal shelves^1, 37^) and E15.5 (the stage of fusion of palatal shelves^1, 37^) wherein osteogenic marker genes were identified in higher abundance than the earlier phase of development (E13.5). Importantly, the inverse patterns of expression between the regulatory transcription factors, *Twist1* and *Twist2*, and the pre-osteoblast orchestrator of osteogenic differentiation, *Runx2*, provide a robust mechanistic framework of osteogenic induction at this stage of palatal fusion. This inverse regulatory relationship between *Twist1/Twist2* and *Runx2* transcriptional control then permits differentiation to continue toward the synthesis of necessary structural components of developing bone (e.g., *Col1a1, Sparc*)^38^. Thus, this combination of regulatory and structural transcriptional activation likely continues the cascade toward terminal differentiation of osteogenic mesenchymal cells in the fusing palate, similar to what is seen mechanistically in the developing teeth^39–41^. Specifically, E15.5 was marked by increased osteoblast differentiation and commitment marker gene abundance, as well as the introduction of marker genes identifying terminally differentiated bone cells (osteocytes) within the hard palate mesenchyme. Osteocytes were found to be spatially restricted in expression lateral to less differentiated osteoblasts and pre-osteoblasts, which had already begun traversing medially toward the mid-palatal suture from their lateral ossification zone of origin, as well as condensed expression enrichment within the midline mesenchyme of the newly fused palate. Importantly, the timeline of transcription for important osteogenic genes identified herein provide valuable molecular insights to corroborate what is known histologically through palatal development (Fig. 7). Explicitly, the presence of abundant osteogenic gene transcripts at E15.5 warrants further studies on the origin of osteogenic cells within the midline of palatal shelves through lineage tracing and live cell imaging modalities as post-fusion development of the mid-palatal suture proceeds.

Given the timeline of osteogenic gene expression observed in our sequencing studies herein, future studies wishing to assess the functional role(s) of osteocytes in differentiating palatal mesenchyme would require studying stages beyond E15.5. This approach is dictated by the markedly low mRNA expression levels of osteocyte-specific markers that were identified in earlier palate developmental stages through bulk, single-cell and spatial RNA-seq modalities. Notably, therapeutic control of osteogenic processes *in vivo* has been achieved and optimized previously with a targeted Wnt pathway agonist (e.g., Romosozumab, monoclonal antibody to sclerostin) to modulate the regulatory function of osteocytes for the bolstering of bone tissue^42–44^. The possibility of transiently manipulating the regulatory signaling control exerted on proliferating and differentiating pre- and functional osteoblast cells by osteocytes may provide exciting avenues of potential therapeutic development for congenital deficiencies of craniofacial bone tissues, as our lab^45, 46^ and others^47, 48^ have demonstrated previously through targeting other up-stream Wnt pathway regulators. As the future of fetal and neonatal medicine includes *in utero* preventive and therapeutic interventions, there is a need to continue to decipher the spatiotemporal gene regulatory networks driving normal and abnormal development to then be able to intervene safely and effectively with targeted therapeutics^10, 49–52^.

An important component of this transcriptomic profiling analysis included the identification of up-regulated marker genes not previously reported in the process of palatal fusion (*Deup1*, *Dynlrb2*, *Lrrc23*). Prior work in palate development has largely focused on known mouse genetic models as the basis of studying genes of relevance to known causative mutations in humans. Now, with the ability to capture entire tissue-specific transcriptomes across space and time, the possibilities for discovery of novel relevant marker genes critical for palatogenesis have expanded significantly. *Deup1*, a deuterosome-mediated centrosome amplifier, has a function of promoting multi-ciliated epithelial cell differentiation^53^. Similarly, dynein light chain roadblock-type 2 (*Dynlrb2*) is known to be associated with microtubule-based cell movement, located in the ciliary tip^54^. Given the multiple gene markers indicative of primary cilia involvement in palatal fusion, further study should investigate the role of specific bone cells (osteoprogenitors, pre-osteoblasts, osteoblasts, and osteocytes) through development and their respective cilia-mediated cell-cell interactions. Leucine-rich repeat-containing 23 *(Lrrc23)* has shown some evidence of interaction with the CD28 protein in the development of regulatory T cells, but its function remains largely unknown^55^. Other Lrr family proteins are known for participating in protein-protein interactions^56^.

Several of the genes listed among the top differentially expressed genes in our spatial transcriptomics analysis are implicated in primary cilia, consistent with prior work linking osteogenic differentiation and cilia function^57^. In fact, ablation of ciliary structural proteins can lead to inhibited osteoblast differentiation, osteoblast polarity, and an overall reduction in bone formation^58^. The pathway analyses from our bulk RNA-seq dataset, corroborated *in situ* with three of the five top DEGs identified in the spRNA-seq differential analysis related to cilia function, highlight the likely functional importance of primary cilia in orchestrating the differentiation of palatal mesenchyme into palatal bone at (and likely after) the stage of fusion. This provides yet another layer of evidence in support of previous studies which have linked conserved primary cilia function and palatogenesis^59–61^. Taken together, there seems to be a strong biologic rationale in focusing on embryonic palatal osteogenesis to further explore the role of primary cilia in this process, and its role in the pathogenesis of craniofacial dysmorphologies.

In summary, this novel spatiotemporal transcriptomic map of secondary palate development provides for the first time a transcriptome-wide, staged developmental atlas of temporally and spatially resolved morphogenetic gradients culminating in palatal fusion. Osteogenic lineage mapping in the secondary palate mesenchyme enabled greater insight to the molecular effectors and cellular populations driving the transitory stage of palatal shelf contact, adhesion, and fusion. Novel enriched genes specific to the palate were identified and mapped spatiotemporally at the site of fusion. Such knowledge will advance understanding of the basic mechanisms involved in the complete fusion of palatal shelves, to be referenced in tandem with preclinical disease model studies of cleft palate in vertebrates to further the discovery of novel candidate genes and innovative therapeutics for cleft palate anomalies in humans.

## METHODS

### Animals

All animal procedures were approved by the National Institutes of Health, National Institute of Child Health and Human Development Animal Care and Use Committee (ACUC), under Animal Study Protocol (ASP) #21-031. C57BL/6J mice were obtained from the Jackson Laboratory. Inbred strain of female C57BL/6 mice were utilized for all experiments. 4-12 weeks-old female mice were used for mating. Healthy fertile male mice were mated with the same strain C57BL/6J female mice. Timed pregnancies were conducted via vaginal plug identification, with day 0.5 indicating date of identification.

### Palate Dissection, Single Cell Dissociation

Pregnant mice were sacrificed by CO_2_ inhalation and cervical dislocation. All surgical procedures were performed by using a surgical loupe (Orascoptic, Eye Zoom, 5.5x magnification). Pregnant mice were placed in the supine position on a sterile, absorbent surgical pad and disinfected with 70% ethanol along the site of planned incision. An incision was made on the abdomen along the midline using small surgical scissors. A fresh microsurgical scissor was then used to carefully incise the peritoneum to expose the uterine chain. Using a blunt forceps, the uterine chain was externalized. Embryos were dissected out by releasing it along the myometrium, incising at the oviduct bilaterally and the median uterine horn ligament attachment. Whole embryos were transferred into ice-cold sterile PBS in a 10 cm petri dish. Each embryo was carefully dissected out from the uterus and extra-embryonic amnion and chorionic tissues, then transferred to a new 10 cm culture dish with fresh ice-cold PBS. A blunt forceps was used to hold each embryo and by using a small fine microsurgical scissor, an incision was made on bilateral oral commissures, allowing for extended opening of the mandible and clear vision of the palate cranially. Careful microdissection of the palatal shelves only from respective embryonic stages was performed, with noted potential extra-palatal tissue contamination due to surgical imprecision. Pooled littermates (n=3 biological replicates per sample) of each respective stage were utilized for either total RNA isolation (E13.5, E14.5, E15.5, E16.5, P0) or single cell dissociation (E13.5, E14.5, and E15.5), following manufacturer protocols (10x Genomics).

### Total RNA Isolation, Bulk RNA-sequencing

Dissected palatal shelves from E13.5, E14.5, E15.5, E16.5, and P0 were used for bulk RNA-sequencing. Secondary palate tissue samples from 3 embryos were pooled into 1.5 mL tubes and placed immediately on dry ice and stored in −80°C. For each stage of analysis, 3 technical replicates and 3 biological replicates were included for sequencing. Tissues were homogenized using biomasher II. mRNA was extracted and purified using Macherey Nagel™ NucleoSpin™ mini kit. 1-4 ug of total RNA samples were purified with PolyA extraction, and then purified mRNAs were constructed to RNA-Seq libraries with specific barcodes using illumina TruSeq Stranded mRNA Library Prep Kit. All the RNA-Seq libraries were pooled together and sequenced using illumina NovaSeq to generate approximately ∼40 million 2×100 paired-end reads for each sample. The raw data were demultiplexed and analyzed further. Raw sequence reads were processed using lcdb-wf v1.9rc (lcdb.github.io/lcdb-wf/) according to the following steps: Raw sequence reads were trimmed with cutadapt v3.4^62^ to remove any adapters while performing light quality trimming with parameters ‘-a AGATCGGAAGAGCACACGTCTGAACTCCAGTCA -A AGATCGGAAGAGCGTCGTGTAGGGAAAGAGTGT -q 20 –minimum-length = 25.’ Sequencing library quality was assessed with fastqc v0.11.9 with default parameters. The presence of common sequencing contaminants was evaluated with fastq_screen v0.14.0 with parameters ‘–subset 100000 –aligner bowtie2.’ Trimmed reads were mapped to the Mus musculus reference genome (GENCODE m18) using HISAT2 v2.2.1.^63^ Multimapping reads were filtered using samtools v1.12.^64^ Uniquely aligned reads were then counted in genes with the featureCounts program of the subread package v2.0.1 using Mus musculus reference (GENCODE m18) annotations.^65^ Differential expression was performed using raw counts provided to DESeq2 v1.34.0^66^ with the following modifications from lfcShrink default parameters: type=“normal” and lfcThreshold=1. A gene was considered differentially expressed if the false discovery rate (FDR) was <0.1 (default for DESeq2 for the statistical test that magnitude of the log2FoldChange is greater than 1 (lfcThreshold=1)). Patterns of expression profile were computed and plotted using the function degPatterns from DEGreport v1.30.0 (http://lpantano.github.io/DEGreport/). Functional enrichment was performed for GO Biological Process, Cellular Component and Molecular Function using the ClusterProfiler v 4.2.0 function go.enrich.^67^

### Single-cell RNA-sequencing (scRNA-seq)

The samples were constructed into scRNA-seq libraries using 10X Chromium Next GEM Single Cell 3’ Reagent Kits, v3.1 following the manufacturer’s instructions. The volume of each single-cell suspension loaded onto the Chromium was adjusted for a targeted cell recovery of 5000-6000. ds-cDNA generation was validated using a High Sensitivity chip on an Agilent 2100 Bioanalyzer. The number of PCR cycles was adjusted based off the concentration of each ds-cDNA sample. Prior to PCR, each ds-cDNA sample was ligated with a unique index adapter. Final scRNA-seq libraries were validated using a High Sensitivity chip on an Agilent 2100 Bioanalyzer. The libraries were then pooled together and sequenced on a Nova Seq 6000 using an SP flow cell. Initial processing and quantification of single-cell gene expression was performed using the 10X Genomics Cell Ranger analysis pipeline against the pre-built mm10 reference genome based off GENCODE vM23/Ensembl 98 (gex-mm10-2020-A). Briefly, each sample was demultiplexed using Cell Ranger mkfastq (default parameters) using specified library barcode mixes then aligned and quantitated using Cell Ranger count (with --include-introns flag set).

### scRNA-sequencing Analysis

Reads were aligned to the mouse genome (mm10) before generating a gene by barcode count matrix using Cell Ranger (10X Genomics) with default parameters. In total, ∼9,400 cells were obtained from the original Cell Ranger pipeline. To analyze high quality cells only, we used scDblFinder package with default parameters to identify and remove doublets from further analysis in each of the datasets. Cells were further filtered based on the percentage of counts associated with mitochondrial mapped reads and number of detected genes per cell, and we removed cells > 3 median absolute deviations (MADs) from the median of the total population. Filtration of cells based on quality control metrics, data normalization, scaling, dimensionality reduction, highly variable feature detections, population sub-setting, data integration and differential expression were all performed with the Seurat v4 package based on the standard gene expression and differential expression workflows. Multiple parameters sets were run for clustering resolutions, with manual inspection of each set of results to identify the most concordant with known palate biology. Final parameters selected were the default Seurat parameters with the following modifications: resolution parameter was 0.2, Wilcoxon algorithm was used for finding markers, and scaling counts was performed after regressing out the mitochondrial transcript and ribosomal transcript percentages, before non-linear dimensional reduction and community detection were performed. Applying these stringent filtration steps resulted in 3488 cells for WT13.5, 1756 for WT14.5 and 2585 for WT15.5. The 3 datasets were integrated after performing the SCTransform normalization method.^68^ Cell-types were assigned post-analysis to clusters after inspection of the marker genes. Differential expression was performed on the following contrasts: WT14.5 vs. WT13.5, WT15.5 vs. WT14.5, WT15.5 vs. WT16.5.

### Fluorescent Multiplex mRNA *In Situ* Hybridization (RNAScope)

Mouse embryos were collected in biological triplicates at E13.5, E14.5, and E15.5 and fixed in 10% formalin for 24 hours. Samples were then processed to paraffin embedding and were sectioned at 5 µm on a microtome. RNAScope multiplex fluorescent v2 assay (Advanced Cell Diagnostics, 323100) was used for *in situ* hybridization according to the manufacturer’s instructions with modified pretreatment custom reagent for antigen retrieval. Positive and negative control probes were employed with assistance from Advanced Cell Diagnostics’ Professional Assay Services to ensure quality and reproducibility of RNA assays in our embryonic mouse FFPE samples. Marker probes from Advanced Cell Diagnostics for *Col1a1*, *Sparc*, *Runx2*, and *Sost*, were used in this study. Representative serial slides were also stained with hematoxylin and eosin (H&E) for histomorphometry context. The fluorescent slides were imaged on an AxioScan.Z1 slide scanner (Zeiss) with Plan-apochromat 40x/0.95 objective in five fluorescent channels (DAPI, FITC, Cy3, Cy5, Cy7).

### Spatial RNA Sequencing (spRNA-seq, 10x Genomics, Visium)

All steps from 10X Genomics’ FFPE Visium Spatial workflow were followed. In brief, spatial RNA sequencing (spRNA-seq) slides are coated with an array of poly-T primers, which encode unique spatial barcodes. These barcodes contain thousands of encoded oligonucleotides within a catchment frame of 6.5 x 6.5 mm. The oligonucleotides are selectively hybridized with the 3’ end of mRNA eluted upon tissue permeabilization, enabling scRNA-seq-like mRNA sequencing with the individual barcode spots replacing the individual suspended cells. Each barcoded spot on the slide is ∼55 µm in diameter, which is predicted to capture ∼10 cells per spot. FFPE tissue was sectioned directly onto the barcoded slide, H&E stained, then underwent enzymatic permeabilization, which allowed for mRNA release and subsequent capture by primer-coated slides. These mRNA molecules are visualized through the incorporation of fluorescent nucleotides into the complementary DNA (cDNA) synthesis process. The resolution of unique spatial barcoding *in situ* allows matching of RNA abundance with the original spatial location in the tissue section, providing a whole-transcriptome RNA-sequencing with 2-dimentional spatial relation. All differential spatial sequencing analyses were conducted using the 10X Genomics Loupe Browser (Space Ranger).

## Acknowledgments

We thank Dr. Vincent Schram of the Microscopy and Imaging Core (NICHD, NIH) for their help with the fluorescence imaging; Drs. Sergey Leiken and Elena Makareeva (NICHD, NIH) for their technical and analytical support in establishing our independent next-generation sequencing workflow and multiplexed *in situ* hybridization assays. We are grateful to Ms. Nichole Swan, Computer Support Services Core (NICHD, NIH) for her help in constructing the schematics, as well BioRender and CleftGeneDB for generic schematics. Finally, we thank Dr. Gerard Karsenty (Columbia University) for his feedback on data visualization and interpretation during manuscript preparation.

## Funding

National Institutes of Health, National Institute of Child Health and Human Development, Intramural Research Training Award (JOP); National Institutes of Health, National Institute of Child Health and Human Development, Intramural Research Program (RDS)

## Author contributions

Conceptualization: JOP, RDS; Methodology: JOP, RDS, RKD, FK; Investigation: JOP, RDS, RR, DMR, PC, FK; FRF, JI, AM, KC, GF, CE, RKD, RDS; Visualization: JOP, RDS, DMR, FRF, JI, AM, KC, GF, CE, RKD; Funding acquisition: JOP, RDS; Project administration: JOP, RKD, RDS; Supervision: RDS; Writing – original draft: JOP; Writing – review & editing: JOP, RR, DMR, RKD, RDS

## Competing interests

Authors declare that they have no competing interests.

## Data and materials availability

All results and analysis data are available in the main text or the supplementary materials. Additional information and materials are available from the corresponding author upon reasonable request. All sequencing data reported in this paper are deposited in GEO under the SuperSeries GSE205449, which includes bulk RNA-seq GSE205446 and Single Cell RNA-seq GSE205448. Further, all sequencing data are deposited on the open-access platform for craniofacial researchers, FaceBase.

## Notes

### Competing Interest Statement

The authors have declared no competing interest.

