## Supplemental File for "Integrated spatiotemporal transcriptomic resolution of embryonic palate osteogenesis"

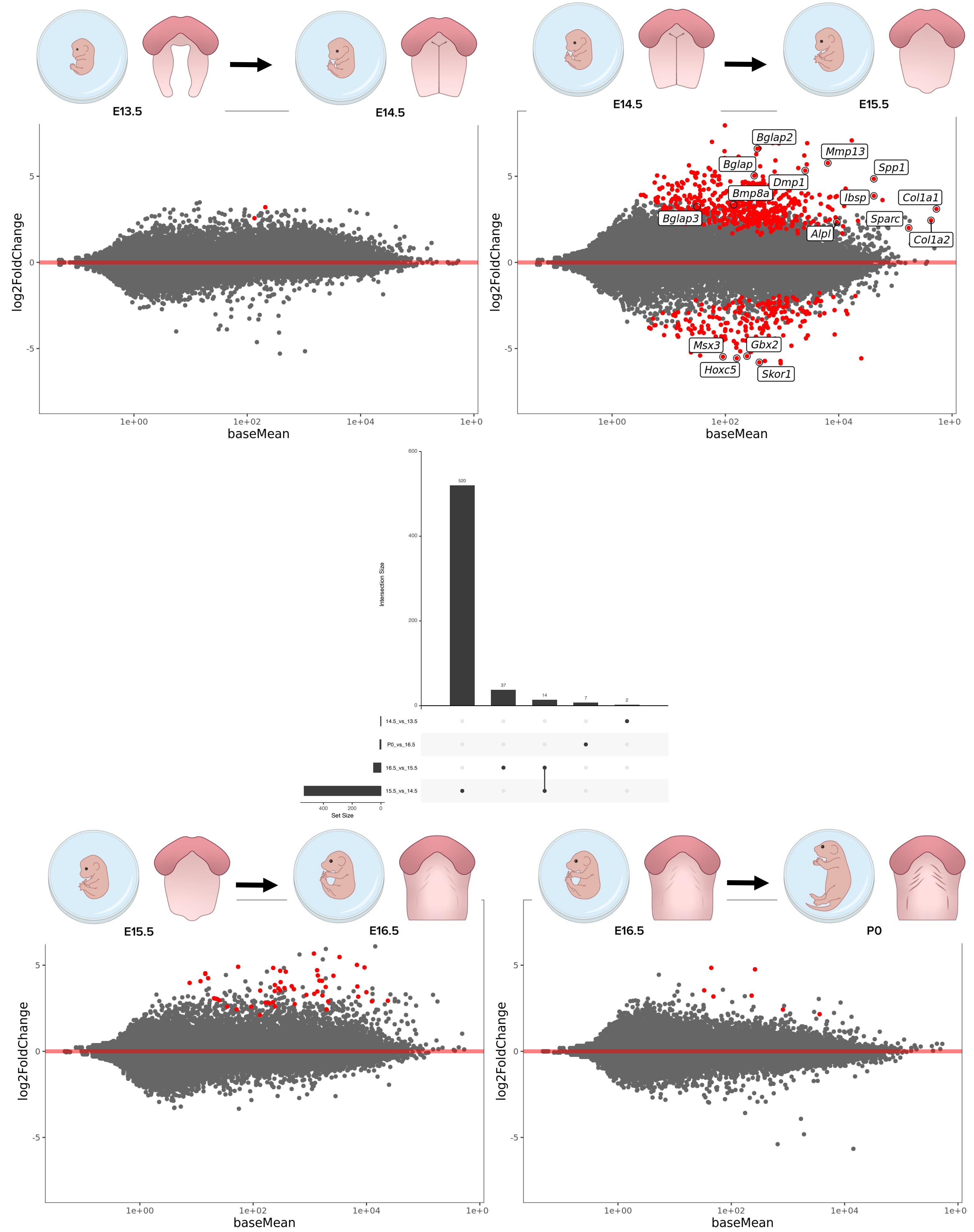


**Supp. Fig. 1. Staged differential expression analyses identify E14.5-E15.5 transition as greatest overall shift in gene expression.** Grouping genes into expression patterns by Z-score (abundance) corroborated the clustered expression patterns pre-fusion and post-fusion identified in heatmap and PCA. Closer investigation of the stage of palatal fusion (E14.5-E15.5) demonstrated the greatest shift in overall gene expression compared to all other staged comparisons.

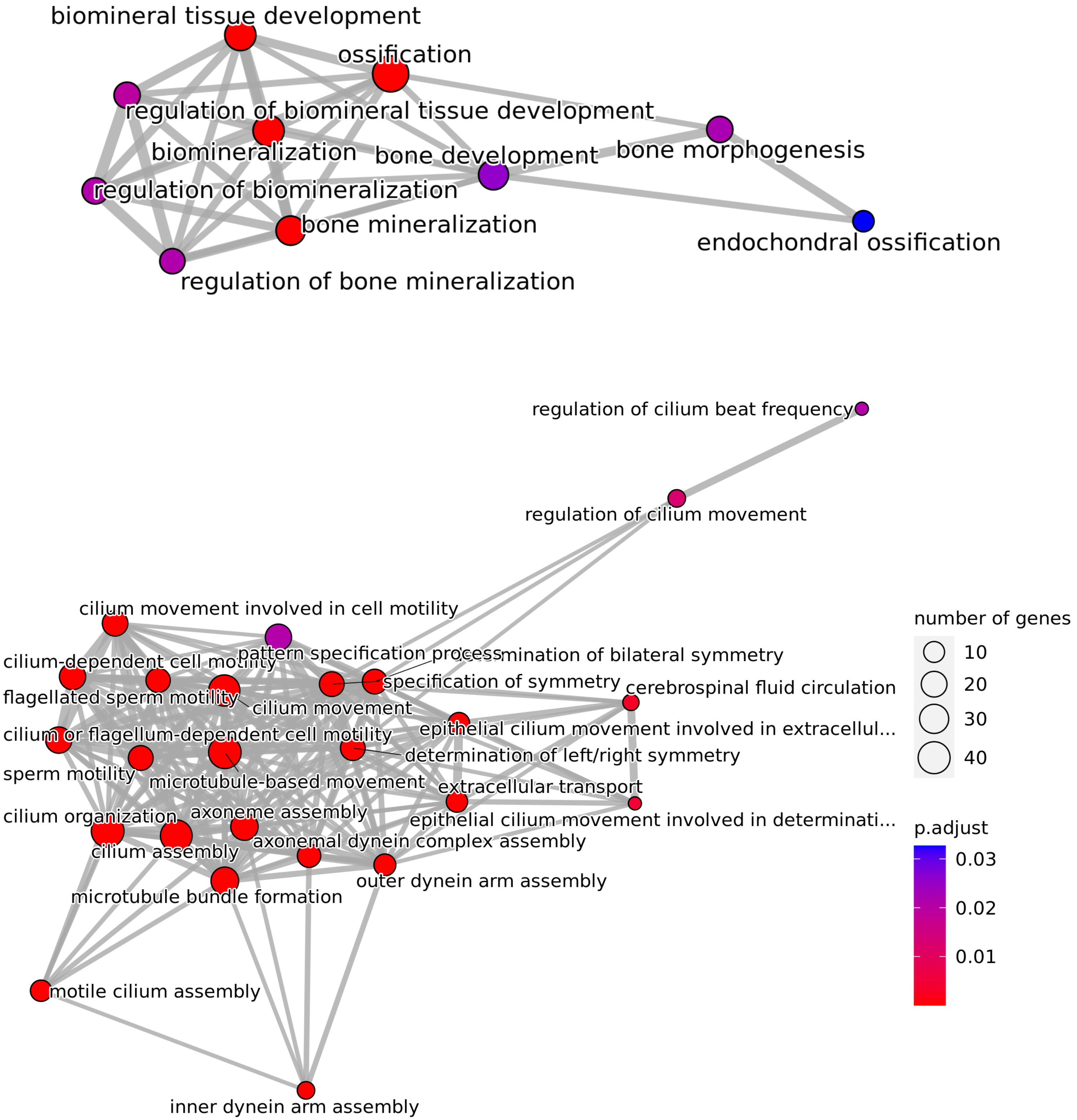


**Supp. Fig. 2. Functional Enrichment Analysis from E14.5 vs E15.5 highlights bone morphogenesis and cilia development as key processes at this timepoint.** Functional enrichment analyses from GO Biological Processes database corroborated gene patterning associations based on stage-specific identified from single genes.

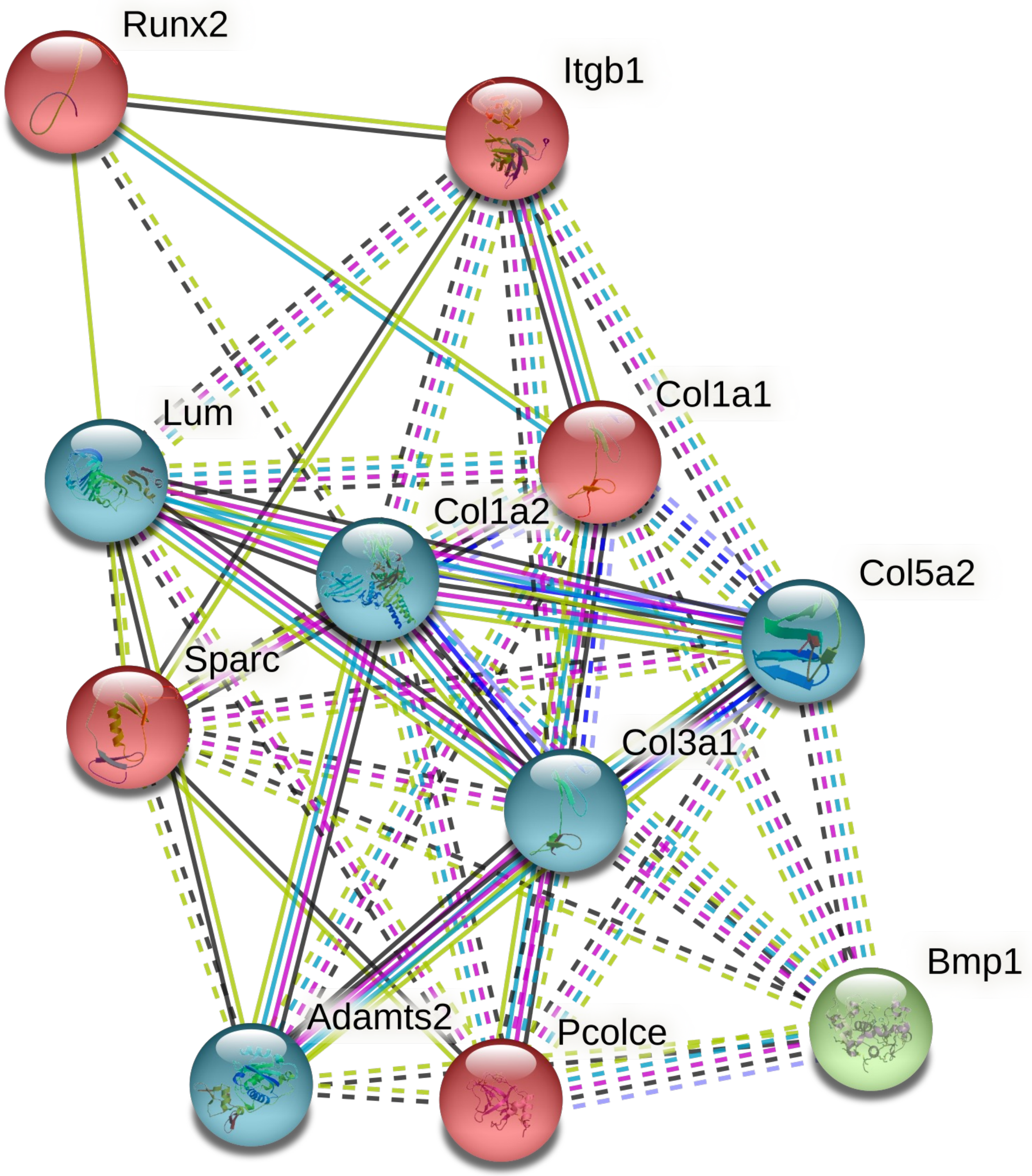


Cluster 1

Tissue Expression Database: **Osteoblast [BTO:0001593]**
*(Col1a1, Sparc, Runx2)*

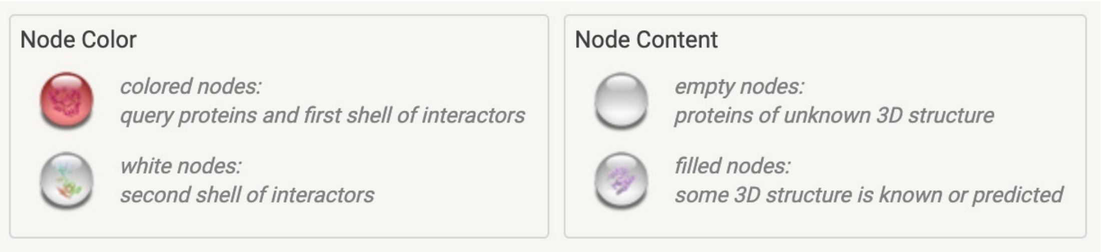


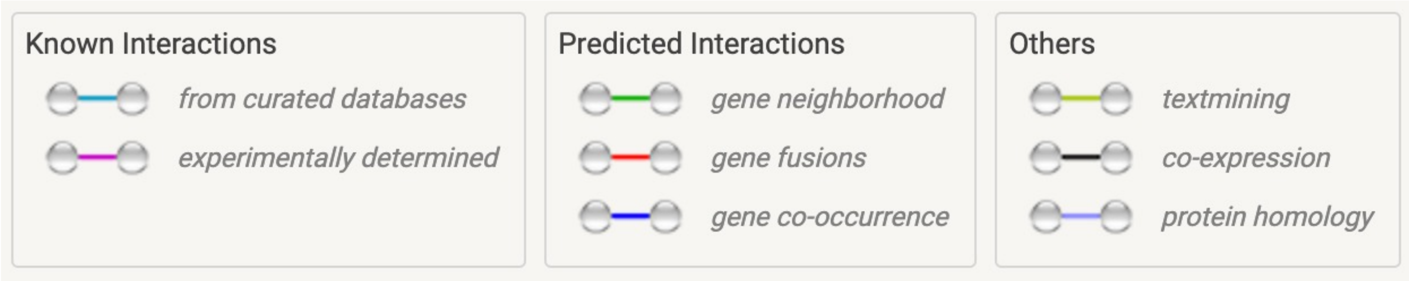


**Supp. Fig. 3. From String Consortium Database, K-means clustering highlights osteoblast markers with common interactions with high strength of association (2.41) and low false discovery rate (1.20e-06).**

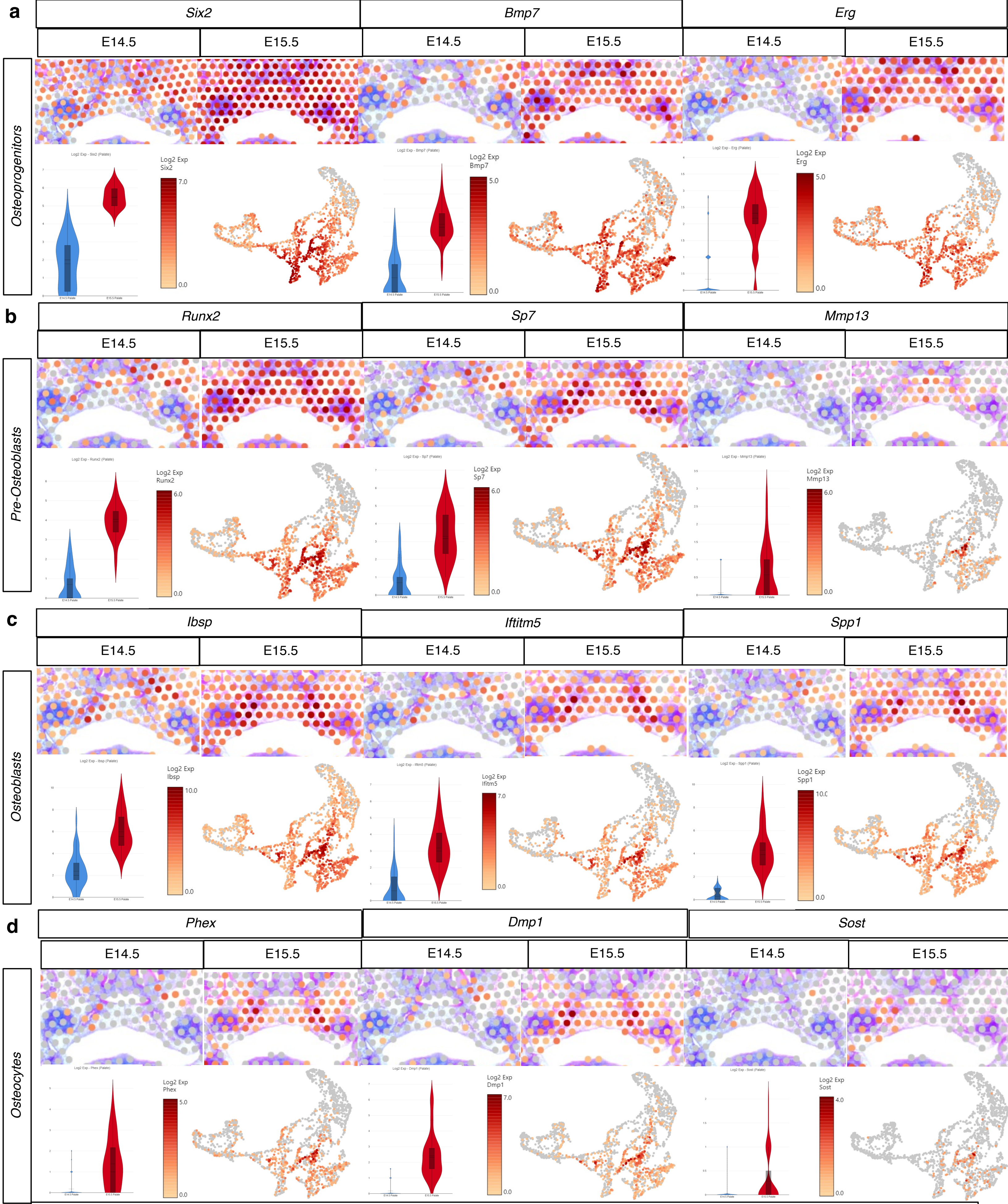


**Supp. Fig. 4**. ***In vivo*** **spRNAseq confirms temporal increase in osteogenic markers and provides spatial quantification of lineage-specific differentiation of condensed mesenchyme progenitors of palatal bone**. Markers of osteogenic lineage induction included the earliest **a** osteogenic progenitors *Six2, Erg*, and *Bmp7*, then **b** pre-osteoblast markers *Runx2*, *Sp7*, and *Mmp13*, followed by **c** more mature, functional osteoblast markers *Ifitm5, Ibsp*, and *Spp1*; finally, **d** markers of osteocyte activity *Sost, Dmp1*, and *Phex*, increase at this stage, yet to a lesser extent than earlier markers. Notably, markers from all early osteogenic cell types (osteoprogenitors, pre-osteoblasts, and osteoblasts) demonstrated some expressing at E14.5; however, all osteocyte conserved markers only appeared in substantial expression by E15.5.

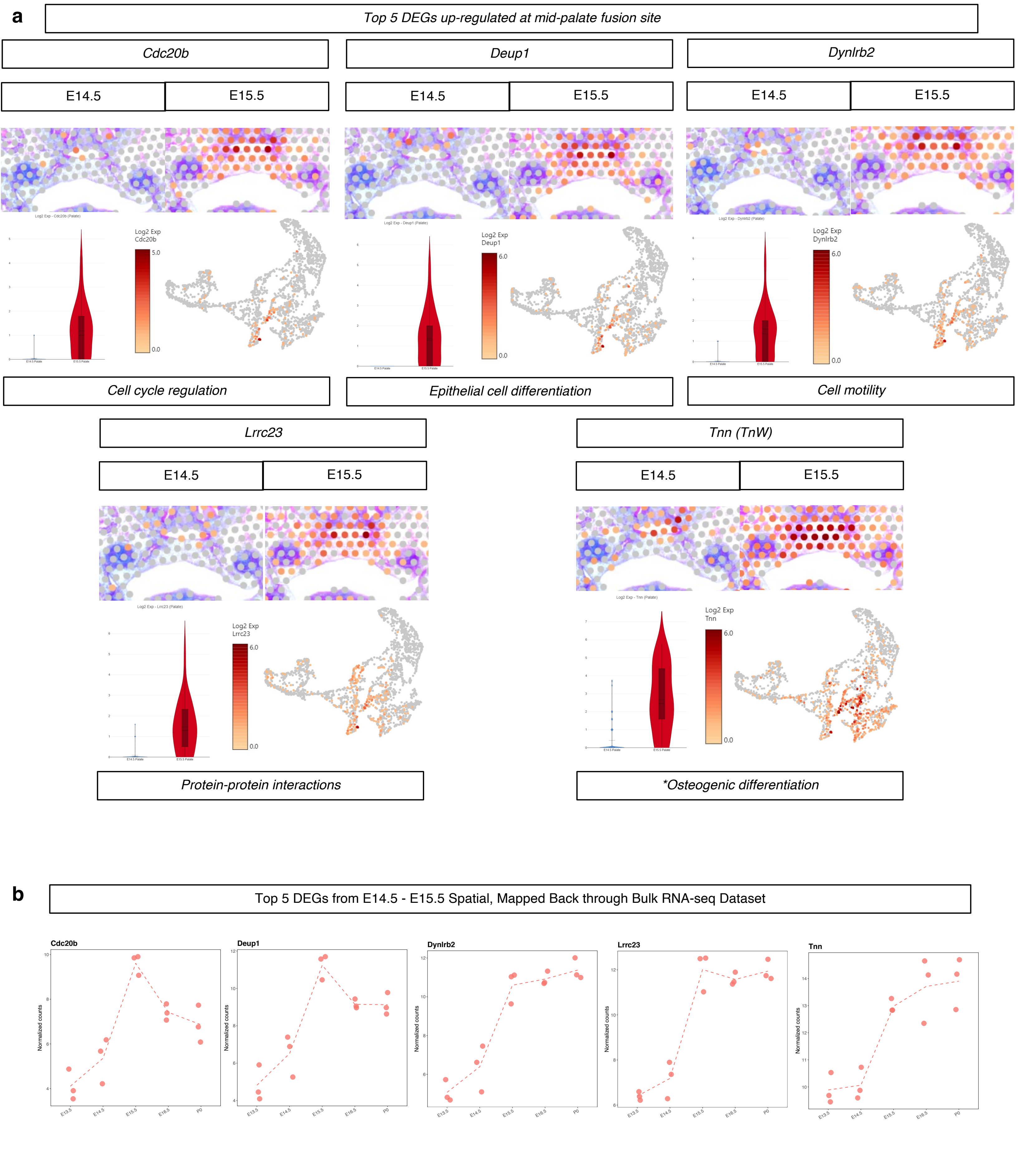


**Supp. Fig. 5. Spatial RNA-seq-derived novel marker genes differentially expressed at the stage of palatal fusion**. **a** Spatial *in vivo* expression, UMAPs, and violin plots from the top 5 marker genes from unbiased differential analysis of selective palate barcoding sequencing capture areas from E14.5 – E15.5 mid-palate coronal sections. **b** Gene plots from previously curated bulk RNA-seq analyses corroborate above spRNA-seq data, further evidence of the potential important functional roles for *Cdc20b*, *Deup1*, *Dynlrb2*, *Lrrc23*, and *Tnn* (aka *Tn-W*) in secondary palate development.
